# Delta-5 elongase knockout reduces DHA and TAG synthesis coupled with an increase of heat sensitivity in a marine diatom

**DOI:** 10.1101/2024.04.19.590303

**Authors:** Junkai Zhu, Shuangqing Li, Weizhong Chen, Xinde Xu, Xiaoping Wang, Xinwei Wang, Jichang Han, Juliette Jouhet, Alberto Amato, Eric Maréchal, Hanhua Hu, Andrew E. Allen, Yangmin Gong, Haibo Jiang

## Abstract

Recent global marine lipidomic analysis reveals a strong relationship in the ocean between temperature and phytoplanktonic abundance of omega-3 long-chain polyunsaturated fatty acids (LC-PUFAs), especially eicosapentaenoic acid (EPA) and docosahexaenoic acid (DHA), which are essential for human nutrition and primarily from phytoplankton in marine food webs. In phytoplanktonic organisms, EPA may play a major role in regulating the phase transition temperature of membranes, while the function of DHA remains to be explored. In the oleaginous diatom *Phaeodactylum tricornutum*, DHA is distributed mainly on extraplastidial phospholipids, which is very different from the EPA enriched in thylakoid lipids. Here, CRISPR/Cas9-mediated knockout of *ptELO5a*, which encodes a delta-5 elongase catalyzing the elongation of EPA to synthesize DHA, led to a substantial interruption of DHA synthesis in *P. tricornutum*. The *ptELO5a* mutants show significant alterations in transcriptome and glycerolipidomes including membrane lipids and triacylglycerols under normal temperature (22°C), and are more sensitive to elevated temperature (28°C) than wild type. We conclude that the PtELO5a-mediated synthesis of small amounts of DHA has indispensable functions in regulating the membrane lipid, and indirectly contributing storage lipid accumulation and maintaining thermomorphogenesis in *P. tricornutum*. This study also highlights the significance of DHA synthesis and lipid composition for environmental adaptation of *P. tricornutum*.

## Introduction

Omega-3 long-chain polyunsaturated fatty acids (LC-PUFAs) are receiving increasing attention due to their important roles in human health (Zhang et al., 2019). Many evidences suggest that docosahexaenoic acid (DHA) contributes to the normal development of visual and neurological systems in infants, and eicosapentaenoic acid (EPA) reduces the incidence of cardiovascular disease in middle-aged and elderly adults (Bazinet and Layé, 2014; Lai et al., 2018). Currently, marine fish and seafood are the primary dietary sources of omega-3 LC-PUFAs. However, they do not possess a complete biosynthetic pathway for omega-3 LC-PUFAs, and these marine organisms can only obtain LC-PUFAs from marine phytoplankton, which are the true producers of these important omega-3 LC-PUFAs (Khozin-Goldberg et al., 2016). Global climate change, specifically ocean warming, is affecting marine phytoplankton communities, which may have a significant impact on the abundance of omega-3 LC-PUFAs in the oceans (Boyce et al., 2010; Tan et al., 2022). A recent analyses of global ocean lipidomes showed a strong negative correlation between the abundance of EPA and temperature, and a weak correlation between DHA and temperature (Holm et al., 2022). The reason behind the different impact of ocean warming on EPA and DHA abundance in marine phytoplankton is still unknown.

Limited research suggests that membrane phospholipids containing omega-3 LC-PUFAs in algal cells may function as shield molecules against exogenous or endogenous oxidative challenges in marine environments (Okuyama et al., 2008; Lupette et al., 2018). In addition, the high content of omega-3 LC-PUFA in membrane lipids can regulate the fluidity and stability of cell membranes in response to temperature changes, thus helping phytoplankton to adapt to low temperatures (Ernst et al., 2016). Although studies on phospholipid models have shown that different compositions of omega-3 LC-PUFAs in phospholipids affect different characteristics of membranes, such as membrane structure and lipid interactions, these have not been adequately investigated in phytoplankton (Sherratt and Mason, 2018; Sherratt et al., 2021). It is indisputable that the proportions of different omega-3 LC-PUFAs varied considerably in most algal species, but it is more common for diatoms to have a very high proportion of EPA and low proportion of DHA (Sayanova et al., 2017; Zulu et al., 2018). This suggests that EPA and DHA may play different roles as membrane lipid constituents in some physiological functions of phytoplankton.

Diatoms are an important group of marine primary producers and major source of omega-3 LC-PUFAs, which play vital roles in the global carbon cycle, climate change regulation, and healthy marine food webs (Field et al., 1998; Zulu et al., 2018; Li et al., 2023). *Phaeodactylum tricornutum*, one of the model species of diatoms, contains high amounts of EPA (30% of the total fatty acids) and trace amounts of DHA, which make it a suitable species to study the metabolic pathway of omega-3 LC-PUFAs (Abida et al., 2015). In the DHA synthesis pathway, delta-5 elongase (ELO5) is the key enzyme, that catalyzes the synthesis of DHA from EPA, and regulates the relative DHA content in the fatty acid profile of *P. tricornutum* (Dolch and Maréchal, 2015).

Expression of heterologous ELO5 from picoalga *Ostreococcus tauri* increased DHA levels in *P. tricornutum*, and much of this over-synthesized DHA was retained in phospholipids, while these alterations in fatty acid composition did not affect the growth of the transformed strains (Hamilton et al., 2015). In comparison, endogenous ELO5 in *P. tricornutum* remains understudied. Two endogenous ELO5s, PtELO5a (protein ID: Phtra3_J9255) and PtELO5b (protein ID: Phatr3_J34485), were annotated as delta-5 elongase in the database of *P. tricornutum*. Our previous study revealed the role of *ptELO5a* in the elongation of EPA through heterologous expression in *Pichia pastoris* (Jiang et al., 2014). However, there is still a lack of direct genetic evidence confirming that PtELO5a is required for the biosynthesis of DHA from EPA in *P. tricornutum*. Meanwhile, the effects of interrupting DHA synthesis on lipid metabolism and physiology in *P. tricornutum* remain unclear.

Over the last decades, successful application of gene editing techniques, such as transcription activator-like effector nucleases (TALENs) and the clustered regularly interspaced short palindromic repeats/CRISPR-associated protein 9 (CRISPR/Cas9) approaches in diatoms, have allowed us to target and stabilize modifications to the diatom genome (Daboussi et al., 2014; Nymark et al., 2016; Moosburner et al., 2020). In this report, *ptELO5a* mutants were constructed using CRISPR/Cas9 gene editing, and the phenotypes of these mutants were analyzed by combining lipidomic and transcriptomic data. We also focused on the association between molecular composition of phospholipids and heat sensitivity according to the phenotype of the mutants under heat stress. This work will improve our understanding of the effects of the DHA synthesis pathway on phospholipid composition and physiology in diatoms.

## Materials and methods

*Phaeodactylum tricornutum* wild-type (WT), *ptELO5a*-overexpression (*ptELO5a*-OE), *ptELO5b*-overexpression (*ptELO5b*-OE), *ptELO5a* mutants and *ptELO5a-*complementation (*ptELO5a-*Com) strains were cultured in sterile artificial seawater enriched in F/2 medium (Guillard, 1975). The *ptELO5a*-OE*, ptELO5b*-OE and *ptELO5a* mutants were additionally supplemented with zeocin with a final concentration of 75 mg L^-1^. The *ptELO5a*-Com strains were additionally supplemented with nourseothricin (NTC) with a final concentration of 200 mg L^-1^. The algal cells were maintained at 22°C under 75 μmol photons m^-2^ s^-1^ light intensity and 12 h light/12 h dark cycle. For the measurements of physiological phenotypes, *P. tricornutum* strains in the exponential growth phase were inoculated into 500 mL medium at an initial concentration of 1×10^4^ cells mL^-1^ with constant aeration. Daily samples were taken for cell number counting, nitrate concentration determination, and Nile red staining fluorescence intensity determination (Collos et al., 1999; Yu et al., 2009). For the heat stress experiment, the incubation temperature was set to 28°C and other experimental conditions remained unchanged.

### Plasmid construction and transformation of *P. tricornutum*

To investigate the subcellular localization of PtELO5a in *P. tricornutum* cells, a fusion protein containing PtELO5a and enhanced green fluorescence protein (eGFP) was designed. The coding sequence of *ptELO5a* was cloned into the multiple cloning sites (MCS) of pPha-CG vector to construct the enhanced green fluorescent protein fusion vector. In addition, the coding sequence of *ptELO5b* was cloned into the MCS of pPha-T1 as the vector for *ptELO5b* over-expression. Approximately 1×10^7^ *P. tricornutum* cells in the exponential growth phase were centrifuged at 3000 g for 10 min, and spread onto each 1.2% agar plate containing F/2 media. The plasmids were coated with tungsten powder and transformed into *P. tricornutum* cells by the Bio-Rad Biolistic PDS-1000/He Particle Delivery System (Bio-Rad, Hercules, California, USA) as the previously described method (Zaslavskaia et al., 2000). The bombarded cells were spread onto 1.2% agar F/2 plates with 75 μg mL-1 zeocin for 2-3 weeks to obtain resistant colonies. The *ptELO5a-GFP* transformed algal strains also serve as *ptELO5a*-OE strains.

To construct the *ptELO5a* mutants, the Cas9 target sites of *ptELO5a* with the PAM signal (NGG) were identified by PhytoCRISP-Ex and six suitable sgRNAs were screened. The plasmid used to the gene editing of *ptELO5a* in *P. tricornutum* was constructed by the insertion of the gRNA expression cassettes into the PBR-CAS9-ShBle vector by the Golden Gate Assembly as described by Moosburner et al. (Moosburner et al., 2020). The resulting Cas9-ShBle:sgRNA episome was introduced into *P. tricornutum* cells using the bacterial conjugation method as described in (Karas et al., 2015). Resistant colonies were selected on F/2 plates supplemented with 75 μg mL^-1^ zeocin. All the plasmids and primer sequences used in this study are listed in Supplemental Table S1 and Table S2, respectively.

### Identification and homozygosis of *ptELO5a* knockout mutants

In order to determine whether the resistant colonies were successfully gene-edited by Cas9, the colonies were randomly selected to prepare cell lysates, and 2 μl of cell lysates were used as specific primers for polymerase chain reaction (PCR) amplification of *ptELO5a* sequence on the genome. Forward primer was set for this knocked out sequence and reverse primer was designed for sequences outside the edited region The PCR products were detected with 1% agarose gel and a portion of the products were selected for Sanger sequencing. Genotypic characterization of colonies using a CRISPR Edit Inference Tool (ICE, Synthego, https://ice.synthego.com). Colonies with 100% indel percentage were selected to determine the sequence of the fragment that was knocked out. The cloned genomic DNA and cDNA were verified by PCR for further determination of knockout results.

### Confocal microscopy

*P. tricornutum* cells containing subcellular localization vector (V_ptELO5a-eGFP_) were cultured to exponential growth phase. Samples were taken to excite GFP fluorescence and chlorophyll fluorescence at 488 nm with the laser scanning confocal microscope LEICA TCS SP8 (Leica, Germany), and GFP fluorescence was observed at emission wavelength 500-520 nm and chlorophyll fluorescence was observed at emission wavelength 625-720 nm. Endoplasmic reticulum (ER)-Tracker Red (Beyotime, Beijing, China) was used to visualize ER with an excitation wavelength of 580 nm and emission by 590-640 nm.

### Transmission electron microscopy

*P. tricornutum* cells were harvested by centrifugation at 3000 g for 15 min, then fixed overnight at 4°C with ten volumes of 4 % (w/v) glutaraldehyde fixative diluted with sterilized seawater. The supernatant was removed and rinsed 3 times with 0.1 M phosphate buffer. The samples were stained with 1% (w/v) osmium tetroxide for 2 h at 4°C in the dark, and washed with 0.1 M phosphate buffer for 3 times, and then dehydrated with increasing concentration of ethanol (30, 50, 70, 90% (v/v) in water) for 10 min and 90% acetone for 10 min. 100% acetone was used to infiltrate the samples for 10 min and repeated for 3 times. Then gradually infiltrated for 1 h using a mixture of Pon 812 Epoxy Resin Monomer and acetone in the ratios of 3:1, 2:1, 1:1 and 1:0, respectively. Finally, the samples were polymerized at 37°C for 12 h and 60°C for 48 h. After ultra-thin sectioning, the sections were stained with uranyl acetate and lead citrate. Images were recorded using the HITACHI H-7650 transmission electron microscopy (Hitachi, Japan) at 80 kV.

### Lipid extraction and analysis

When cultured to the stationary phase (7 d, 8 d, and 9 d), 5×10^7^ cells/sample were collected and the total lipids of fresh algal cells were extracted with chloroform/methanol (2:1, by volume) and redissolved with chloroform after nitrogen blowing. The amounts of triacylglycerols (TAGs) were evaluated by thin layer chromatography (TLC) using hexane/diethyl ether/acetic acid (70:30:1, by volume) and visualized with 0.01% (w/v) primuline reagent. LC-MS was performed on the isolated TAGs to obtain TAG profiles as previously described (Xie et al., 2020). For detailed profiling of all glycerolipids in WT and *ptELO5a* mutants, lipids were extracted and fractionated by 1D and 2D TLC, and purified lipid classes were used for assays by LC-MS/MS as described previously (Jouhet et al., 2017) .

### Measurements of photosynthetic parameters

The chlorophyll fluorescence parameters of *P. tricornutum* were evaluated by a pulse amplitude modulation (PAM) fluorometer (AquaPen AP110-C, Czech Republic). To determine the parameters of maximum photochemical efficiency of photosystem Ⅱ (Fv/Fm) and nonphotochemical exciton quenching (NPQ), the samples were dark-adapted for 20 min, and analyzed in a 1 cm cuvette. The rapid light curve (RLC) was obtained using a pre-programmed light curve (LC3) scheme for relative electron transport rate (rETR) analysis. 1×10^7^ algal cells were collected per sample for chlorophyll *a* content determination. The cells were suspended by adding 4 mL of 90% acetone solution and extracted overnight at 4°C under dark conditions. The samples were centrifuged at 5000 g for 10 min at 4℃ and the absorbance values at 630 and 664 nm of the supernatant were measured. The chlorophyll *a* content was calculated based on Chl *a* = 11.47 A_664_ - 0.40 A_630_ (Wright et al., 2005).

### Reactive oxygen species and malondialdehyde determination

Reactive oxygen species (ROS), malondialdehyde (MDA) and total protein quantitative assay kits were purchased from Nanjing Jiancheng Bioengineering Institute (Nanjing, China). Algae samples with a total cell number of 2 × 10^6^ were collected and incubated in the dark for 20 min with the addition of the DCFH-DA probe, then the excess probe was washed away with sterile f/2 medium and the fluorescence values were measured at 488 nm excitation wavelength and 525 nm emission wavelength. 40 ml for each algae sample was extracted and assayed for MDA and total protein content according to the manufacturer’s instructions. The result of MDA content was calculated as nmol per milligram of total protein (nmol mg protein^-1^).

### Transcriptome and RNA-sequencing

The WT and *ptELO5a* mutant cells were harvested at different culture conditions and time points for transcriptome analysis. Under normal culture condition of 22°C, cells were harvested at nutrient replete phase (4 d) and nitrogen depletion phase (8 d), named as exponential phase (exp) and stationary phase (sta), respectively. In heat stress experiment, algal cells at exponential phase were adjusted to a cell concentration of 1×10^6^ mL^-1^ and treated at 28°C on 1 d and 3 d; these cells were named as heat treat 1d (HT1d) and heat treat 3d (HT3d) respectively. Total RNA of algal cells was extracted using TRIzol reagent (Invitrogen, USA) and sequencing was performed on the BGI PE150 platform (Wuhan Generead Biotechnologies Co. Ltd., China). Clean reads were accurately compared with the reference genome using HISAT2 (2.2.1) and quantified using featureCounts (v2.0.1) (Kim et al., 2019).

### RNA extraction and quantitative real-time-PCR

Total RNA was extracted from *P. tricornutum* cells in the exponential growth phase. Cells were harvested from 40 ml of culture medium and centrifuged at 4000 g for 10 min at 4°C to discard the supernatant, and the cells were frozen and ground in liquid nitrogen. RNA was extracted according to the RNeasy MinElute Cleanup Kit operating manual (Qiagen Inc. Germany). Total RNA (1 μg) was reverse transcribed into cDNA using PrimeScript RT reagent kit (Takara Bio, Japan), and expression was quantified using the TB Green^®^ Premix Ex Taq™ II (Takara). Primer sequences used for qPCR are listed in Supplemental Table S2.

### Construction of *P. tricornutum ELO5* complementary strain

The coding region of *ptELO5a* was amplified by PCR using primer pair of ptELO5aKO_com-F/R. Using In-Fusion Cloning, PCR product was ligated to the *EcoR*I/*Sal*I sites of pPha-Cp1, a modified vector to replace the zeocin resistance cassette with a NTC resistance cassette (Slattery et al., 2018). To avoid the continuous editing of complementary *ptELO5a* fragment by knockout-plasmid, *ptELO5a* mutants with knockout-plasmid loss were screened through five rounds of continuous passage of zeocin free F/2 medium. Then, at exponential-phase, the screened *ptELO5a* mutant cells were complemented by microparticle bombardment with the NTC-resistance plasmid pPha-Cp1 containing the WT *ptELO5a* sequence. Selection of algal transformants was done using F/2 agar plates supplemented with 200 μg mL^-1^ NTC.

### Phylogenetic analysis

The ELO sequences were obtained from NCBI (https://www.ncbi.nlm.nih.gov/protein/) (Supplemental Dataset S1). Multiple protein sequence alignment using software MEGA 11 with the embedded ClustalW program. The phylogenetic tree was reconstructed with default setting using the neighbor-joining method.

### Analysis of the global distribution and environmental correlation of ELO5 genes

The abundance of ELO5 and ELO5-like genes in the ocean was analyzed on both the Marine Atlas of *Tara Oceans* Unigenes and eukaryotes metatranscriptomes (MATOUv1+metaT) (Vernette et al., 2022). The geographical distribution of the phytoplankton ELO5 genes based on the *Tara Oceans* dataset was visualized using the online tool (Chiplot). The taxonomic groups of the phytoplankton enriched at each site were counted, and the environmental conditions at the corresponding sites were evaluated. To include as much phytoplankton as possible, the phytoplankton ELO5 genes from 0.8-5 µm, 5-20 µm and 20-180 µm surface layers of the ocean were selected for further analysis in this study. The following nine environmental parameters were chosen for correlation analysis: temperature (℃), salinity (PSU), photosynthetic active radiation (PAR, mol quanta/m**2/day), iron_5m* (µmol/l), ammonium_5m* (µmol/L), nitrite_5m* (µmol/L), nitrate_5m* (µmol/L), PO4 (µmol/L) and Si (µmol/L). Partial Mantel correlations between ELO5 mRNA abundance and environmental parameters were computed using the vegan R software package.

## Results

### Widespread distribution and environmental responsiveness of ELO5 in marine eukaryotic phytoplankton

To determine the possible ecological and physiological functions of ELO5, *Tara Oceans* unigenes and metatranscriptomes datasets were used to analyze the main regional distribution and abundance of ELO5. The results indicated that ELO5 was widely distributed in the global oceans and found at all *Tara Ocean* stations (Fig. 1A). A total of 79204 ELO5 homologs were hit. Phytoplankton accounted for 46% of the total ELO5 homologs, with Dinophyta (16.6%), Chlorophyta (7.9%) and Bacillariophyta (5.9%) being the three major eukaryotic groups of phytoplankton in the ocean (Fig. 1B). To further explore the specific environmental drivers regulating the expression and distribution of ELO5, the distance-corrected dissimilarities in the abundance of ELO5 transcripts in marine eukaryotic phytoplankton were analyzed with respect to the environmental factors. Overall, three environmental factors, i.e., temperature, nitrate and phosphate, were found to be significantly correlated with the abundance of ELO5 (Fig. 1C). Among these factors, temperature was the main factor driving the global distribution of diatoms.

**Figure 1.**
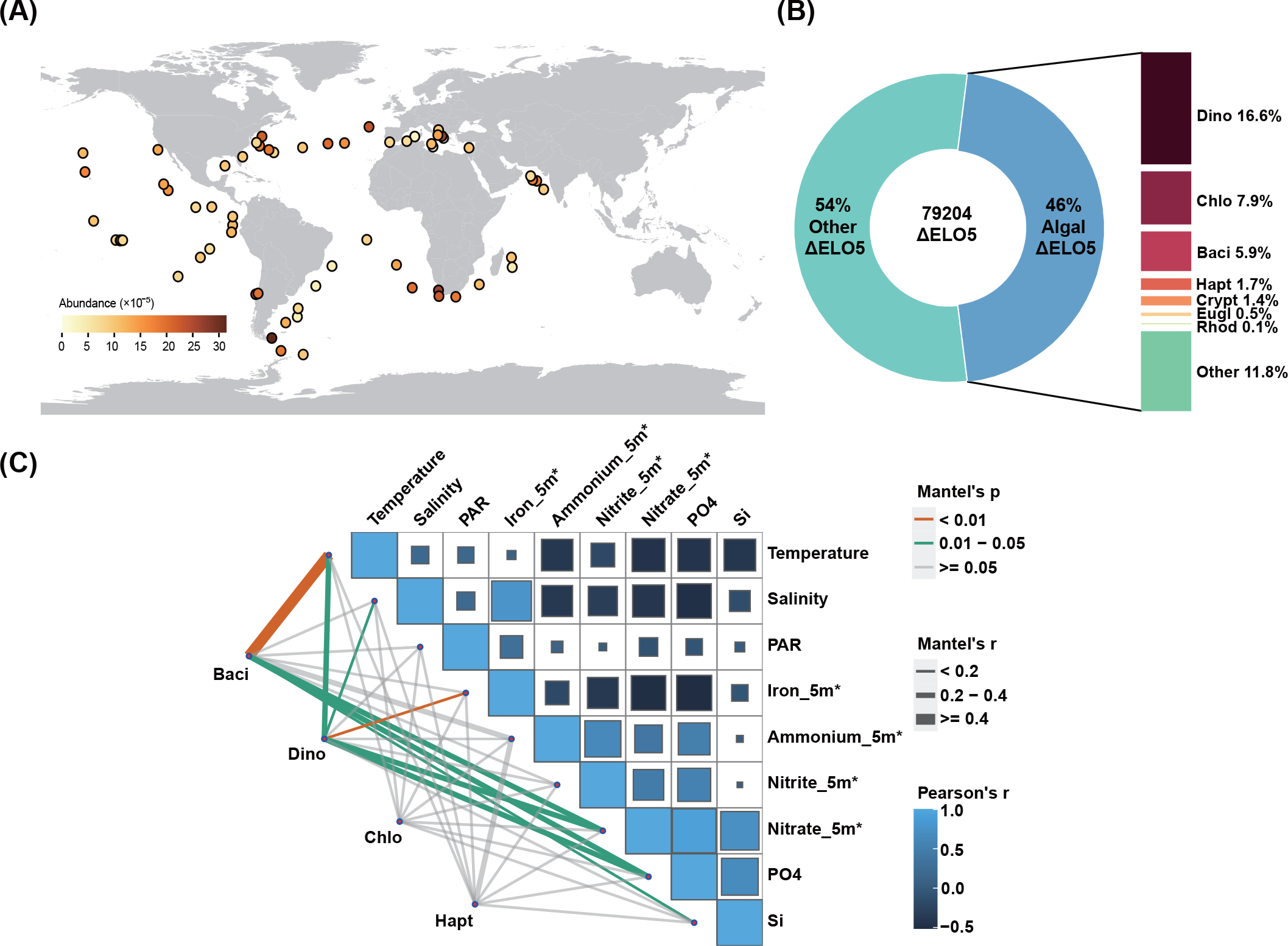
The widespread presence environmental nutrient responsiveness of ELO5 in global marine phytoplankton. (A) Wide geographic distribution of ELO5 found in Tara Oceans. Color scale depicts the abundance of ELO5 mRNA at each site. (B) The wide taxonomic distribution of ELO5 in marine phytoplankton. Dino, Dinophyta; Chlo, Chlorophyta; Baci, Bacillariophyta; Hapt, Haptophyta; Cryp, Cryptophyta; Eugl, Euglenozoa; Rhod, Rhodophyta. (C) Environmental nutrient drivers of phytoplankton ELO5 abundance. Pairwise comparisons of environmental conditions are represented by a color gradient indicating the Pearson’s correlation coefficient. Line width represents the corresponding Mantel’s r statistic for the correlation of taxonomic ELO5 abundance and each environmental factor.

### Bioinformatic analysis, subcellular location, and overexpression of PtELO5

PtELO5a (Phatr3_J9255) and hypothesized PtELO5b (Phatr3_J34485) in *P. tricornutum* contain 369 and 286 amino acid residues, respectively, and are both hypothesized to be membrane-bound proteins, containing ELO-conserved structural domains in their sequences (Fig. S1 and S2). To understand the evolutionary position of PtELO5, phylogenetic analyses were performed using different functional elongases of different organisms (Fig. 2A). The elongases that elongate the carbon chains of long-chain polyunsaturated fatty acids are divided into three main groups. The Δ9 elongases catalyze specific C18 PUFAs, including C18:2Δ9,12 and C18:3Δ9,12,15. Other C18 PUFAs are catalyzed by Δ6 elongases include C18:3Δ6,9,12 and C18:4Δ6,9,12,15. PtELO5a belonged to Δ5 elongases, which are involved in the elongation of C20:4Δ5,8,11,14 and C20:5Δ5,8,11,14,17. FsELO5 from *Fistulifera solaris* showed the highest similarity to PtELO5a in Bacillariophyta, whereas there was a large evolutionary distance between PtELO5b and other typical Δ5 elongases of diatoms. The hypothesized PtELO5b and those elongases with the highest sequence similarity to PtELO5b were clustered into Δ9 elongases.

**Figure 2.**
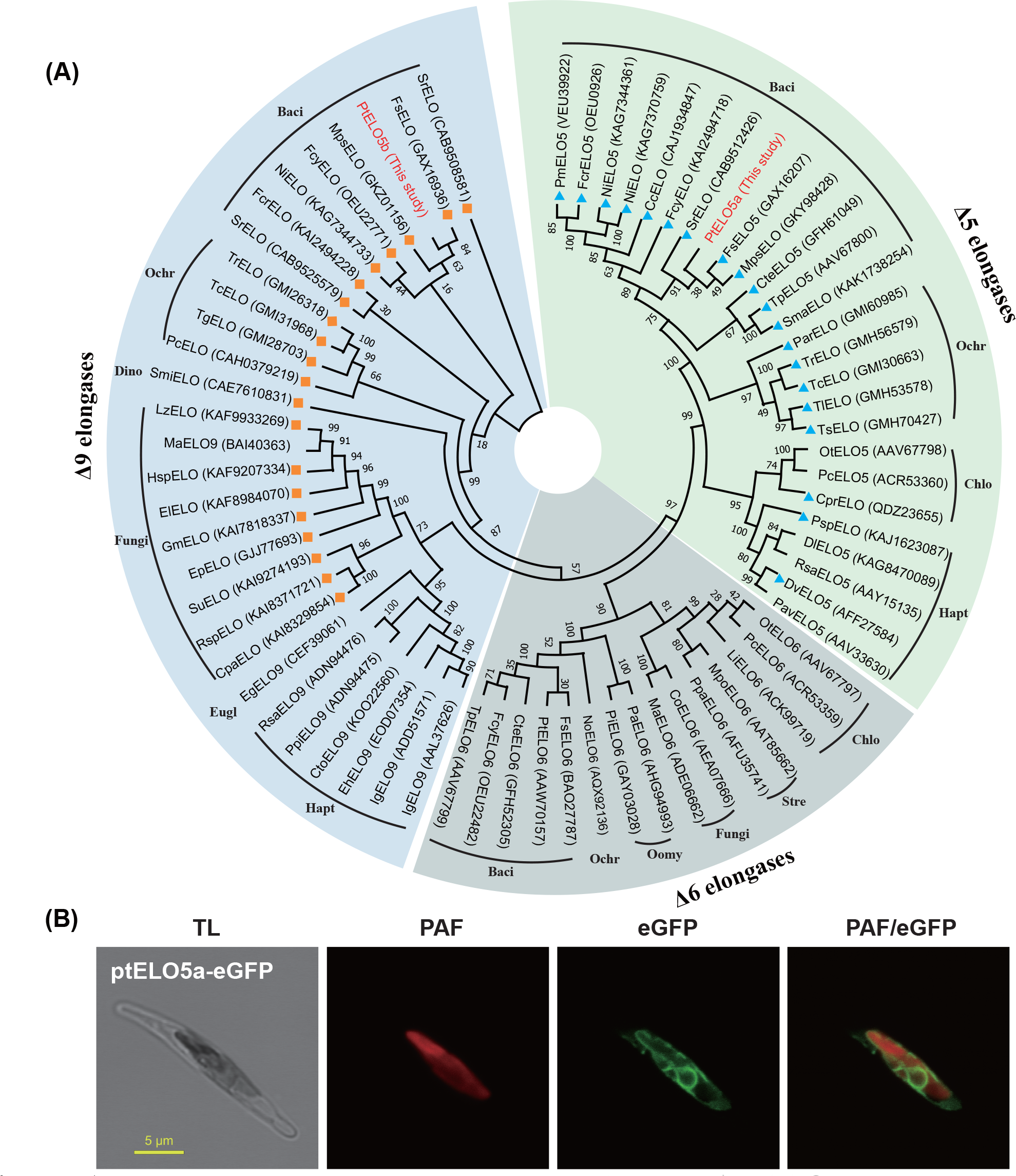
Analysis of PtELO5 sequence alignment and protein localization. (A) Cladogram of ELO5 of difference functions from various organisms. The GenBank ID of the corresponding species ELO5 is shown in bracket. Cc, *Cylindrotheca closterium*; Co, *Conidiobolus obscurus*; Cpa, *Chlamydoabsidia padenii*; Cpr, *Chloropicon primus*; Cte, *Chaetoceros tenuissimus*; Cto, *Chrysochromulina tobinii*; Dl, *Diacronema lutheri*; Dv, *Diacronema viridis*; Eg, *Euglena gracilis*; Eh, *Emiliania huxleyi*; El, *Entomortierella lignicola*; Ep, *Entomortierella parvispora*; Fcr, *Fragilaria crotonensis*; Fcy, *Fragilariopsis cylindrus*; Fs, *Fistulifera solaris*; Gm, *Gamsiella multidivaricata*; Hsp, *Haplosporangium* sp.; Ig, *Isochrysis galbana*; Li, *Lobosphaera incisa*; Lz, *Linnemannia zychae*; Ma, *Mortierella alpina*; Mpo, *Marchantia polymorpha*; Mps, *Mayamaea pseudoterrestris*; Ng, *Nannochloropsis gaditana*; Ni, *Nitzschia inconspicua*; No, *Nannochloropsis oceanica*; Ns, *Nannochloropsis salina*; Ot, *Ostreococcus tauri*; Pa, *Pythium aphanidermatum*; Pc, *Pyramimonas cordata*; Par, *Parmales* sp.; Pav, *Pavlova* sp.; Pi, *Pythium insidiosum*; Pm, *Pseudo-nitzschia multistriata*; Ppa, *Physcomitrium patens*; Ppi, *Pavlova pinguis*; Psp, *Pavlovales* sp.; Pt, *Phaeodactylum tricornutum*; Rsa, *Rebecca salina*; Rsp, *Radiomyces spectabilis*; Sma, *Skeletonema marinoi*; Smi, *Symbiodinium microadriaticum*; Sr, *Seminavis robusta*; Su, *Sporodiniella umbellate*; Tc, *Triparma columacea*; Tg, *Tetraparma gracilis*; Tl, *Triparma laevis*; Tp, *Thalassiosira pseudonana*; Tr, *Triparma retinervis*; Ts, *Triparma strigata*. Baci, Bacillariophyta; Chlo, Chlorophyta;, Dino, Dinophyta; Eugl, Euglenozoa; Hapt, Haptophyta; Ochr Ochrophyta; Oomy, Oomycota; Stre, Streptophyta. The triangular and square markers represent the top 20 sequences most similar to the PtELO5a and PtELO5b protein sequences compared in NCBI, respectively. (B) Subcellular localization of PtELO5a in *P. tricornutum* cells. TL, transmitted light; PAF, plastid autofluorescence; eGFP, enhanced green fluorescence protein; PAF/eGFP, overlay of plastid and eGFP fluorescence.

To verify the subcellular localization of PtELO5a, transgenic *P. tricornutum* cells with over-expressed PtELO5a fused with an eGFP protein were observed by confocal fluorescence microscopy. The green GFP signal was observed around the outermost layer of the plastid and accompanied by partial regional expansion (Fig. 2B). Further staining by ER-tracker was performed to demonstrate the overlap of cER and ER, as cER is a continuous component of the entire ER (Fig. S3). In diatoms, the outermost membrane of the plastid is attached to the ER, known as the ‘chloroplast ER’ (cER) (Gibbs, 1979). Thus, PtELO5a protein is probably located in cER of *P. tricornutum*.

To confirm that *ptELO5a,* rather than hypothesized *ptELO5b,* is essentially required by *P. tricornutum* for the biosynthesis of DHA (22:6), *ptELO5a* and *ptELO5b* overexpressing strains were constructed, respectively (Fig. S4). The analysis of fatty acid composition showed that the relative content of 22:6 in the *ptELO5a* overexpressing strain was increased by 3.67-fold compared with that in WT (Fig. S5). However, the overexpression of *ptELO5b* had no significant effect on the relative composition of 20:5 and 22:6. These results suggest that *ptELO5a* is the main functional gene responsible for carbon chain elongation of 20:5 and the biosynthesis of 22:6 in *P. tricornutum*.

### Identification of the *ptELO5a* knockout mutant obtained through CRISPR/Cas9-mediated gene editing

The *ptELO5a* mutants of *P. tricornutum* was successfully obtained by bacterial conjugation with a Cas9-sgRNA episome-based construct that carried three sgRNA expression cassettes to target three screened loci within the gene. PCR results showed that using WT genomes and total complementary DNAs (cDNAs) as PCR template can amplify the products, which cannot be achieved by using mutant genomes and cDNAs as PCR template (Fig. 3, A and B). DNA sequencing showed that a 35 bp size fragment was missing in the DNA sequence of mutants, compared to the WT DNA sequence (Fig. 3C). These results indicated that the *P. tricornutum* mutants with complete knockout of *ptELO5a* were successfully obtained. Further phenotypic analysis was performed using these three independent *ptELO5a* mutants.

**Figure 3.**
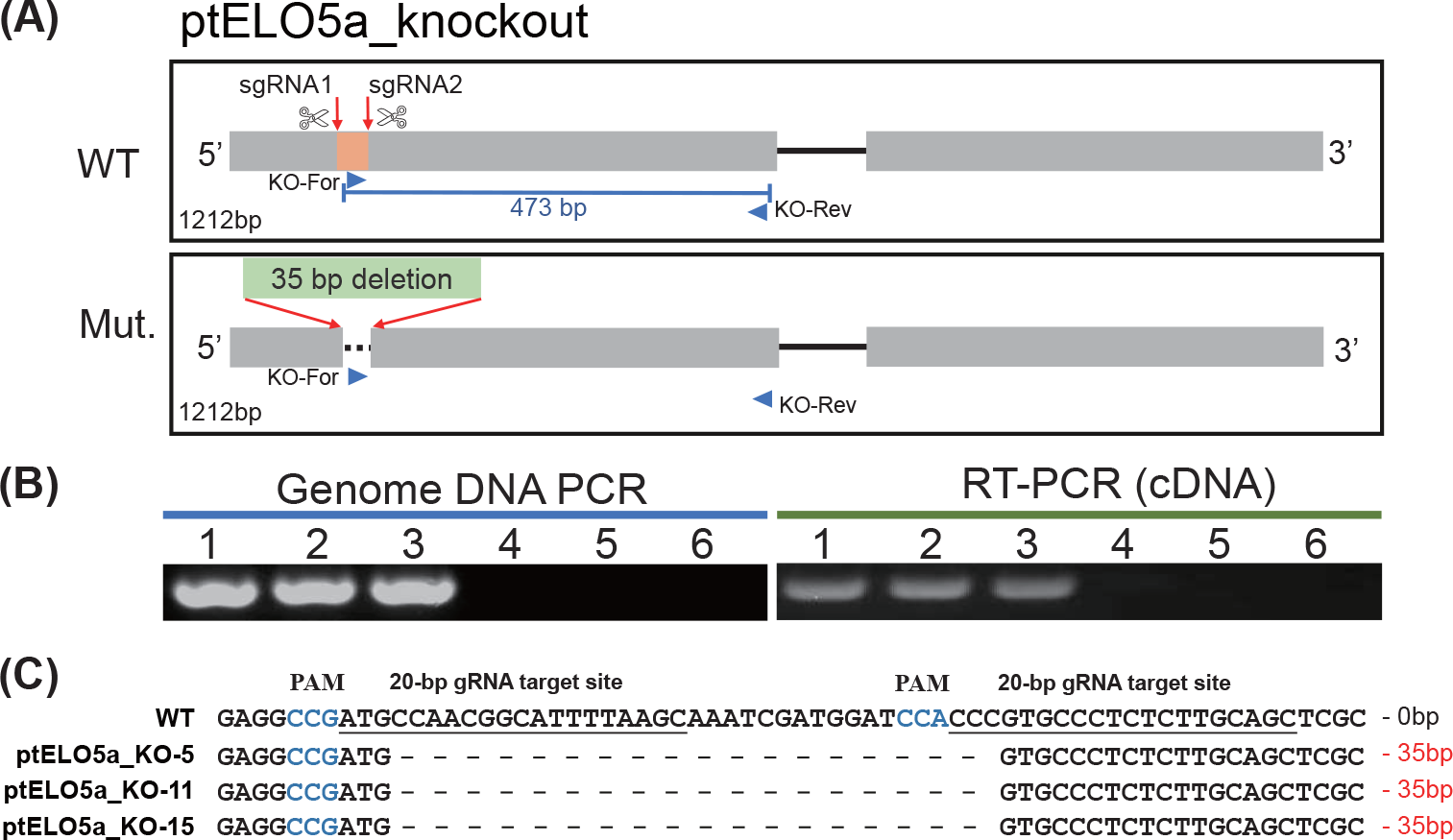
Construction of *ptELO5a* mutant strains. (A) Schematic representations show the fragmental deletion between two targeted sites of the *ptELO5a* mutants; (B) A pair of primers was designed to detect large deletions in the *ptELO5a* mutants by polymerase chain reaction (PCR), using genomic DNAs and total complementary DNAs (cDNAs) as templates. Columns 1, 2, and 3 used the wild type as a template, and columns 4, 5 and 6 used the mutants as a template, respectively; (C) The sequencing results of the successful editing of the mutant *ptELO5a* gene.

### The *ptELO5a* mutants showed increased sensitivity to heat stress

Unlike the consistent phenotype under normal conditions (at 22℃) with sufficient nutrients, the cell morphology of *ptELO5a* mutants appeared significantly different from that of the WT under heat stress conditions (at 28℃), with circular protrusions at both ends of the mutant cells and poor integrity of endomembrane system (Fig. 4, A and B). This subpopulation could maintain a proportion of about 45% in subsequent stress in the mutants (Fig. S6). The mutants almost stopped growing under heat stress, while the WT could still grow slowly at 28℃ (Fig. 4, C and D). In addition, the relative electron transport rate (rETR), maximum quantum yield of photosystem II (F_v_/F_m_), and chlorophyll *a* content in the mutants were all significantly lower than those in WT (Fig. 4, E, F and H). Whereas non-photochemical quenching (NPQ), reactive oxygen species (ROS) and malondialdehyde (MDA) contents in the mutants were significantly higher than those in WT at 28℃, suggesting that the *ptELO5a* mutants were subjected to higher oxidative stress (Fig. 4, G, I and J). Further analysis of fatty acid composition showed that the reduction in PUFA content in the mutants under heat stress was significantly higher than that in WT (Fig. 4, K, L and M). These results indicated that the *ptELO5a* mutants suffered more damage under heat stress than WT.

**Figure 4.**
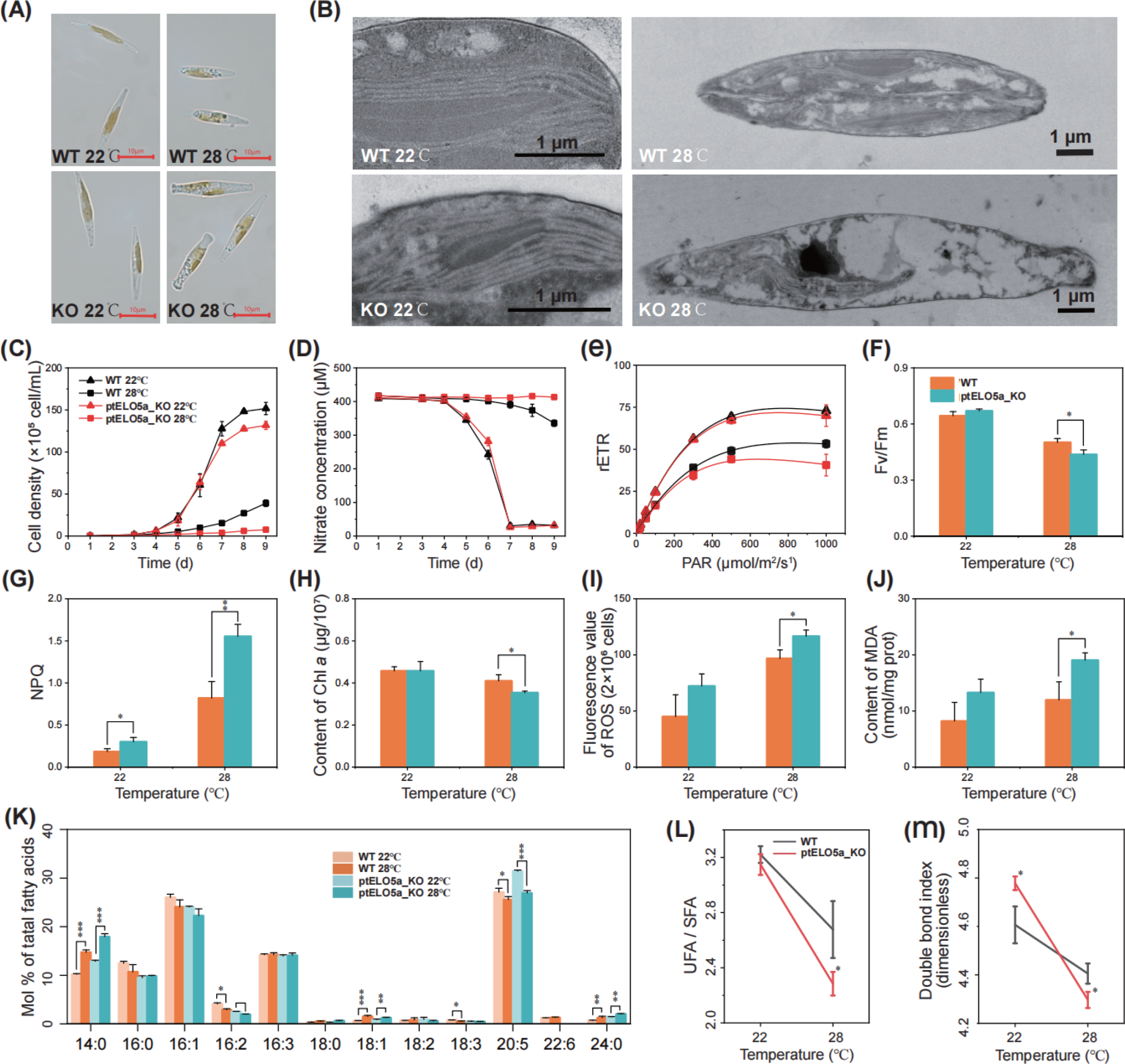
Analysis of the physiological parameters of WT and *ptELO5a* mutants under heat stress. (A) optical microscope observation of morphology; (B) transmission electron micrographs of ultrastructure. (C) cell growth curves; (D) nitrate concentrations in culture medium; (E) relative electron transport rate (rETR); (F) the maximum quantum yield of photosystem II (Fv/Fm); (G) non-photochemical quenching (NPQ); (H) chlorophyll a content; (I) reactive oxygen species (ROS) content; (J) malondialdehyde (MDA) content; (K) Relative composition of fatty acids; (L) Ratio of UFA to SFA. UFA, unsaturated fatty acid; SFA, saturated fatty acids; (M) The double bond index (DBI). DBI = 2 [(% monoenes) + (2 × % dienes) + (3 × % trienes) + (4 × % tetraenes) + (5 × % pentaenes) + (6 × % hexaenes)]/100 is according to Feijão et al., (Feijão et al., 2018). The initial inoculation density was 104 cells mL-1. All 22°C samples were collected at the exponential phase (4th d) and heat-stressed samples were continued to be transferred to 28°C for 3rd d. Data are the average of three biological replicates with error bars indicating standard deviations (n = 3) and the asterisk indicates the significant difference asterisk indicates the significant difference (Student’s t-test, *, *P* < 0.05; **, *P* < 0.01; ***, *P* < 0.001) between the WT and *ptELO5a* mutants.

### Analysis of transcriptome of the *ptELO5a* mutants and the WT under heat stress

After exposure to heat stress at 28℃ for 1 and 3 days, both WT and *ptELO5a* mutants exhibited similar trends in transcript levels across various metabolism pathways. Down-regulations were observed in the ribosome, photosynthesis, protein processing, and fatty acid synthesis pathways, while DNA replication, homologous recombination, mismatch repair, and N-glycosyl synthesis pathways were up-regulated (Fig. S7). The number of significant differentially expressed genes (DEGs) between the mutants and WT increased significantly with increasing duration of heat stress (Supplementary Table S3). Compared to WT, the *ptELO5a* mutants showed a higher degree of inhibition of metabolic pathways associated with protein synthesis and degradation (ribosome, ER protein processing, proteasome, and RNA transport pathways) and photosynthesis (porphyrin and chlorophyll metabolism, 2-oxocarboxylic acid metabolism, and carotenoid biosynthesis) during heat stress. On the contrary, the mutants demonstrated increased activities within various metabolic pathways involved in energy production, notably the citrate cycle (TCA cycle) (Fig. 5, A and B). Moreover, a substantial down-regulation of heat shock protein (HSP)-encoding genes was observed in the mutants compared with the WT under heat stress (Fig. S7). Altogether, these transcriptional results suggested that the knockout of *ptELO5a* interrupted the ER regulation of heat stress, primarily by influencing the coordinated regulation of HSP genes and ER-associated genes involved in the clearance of proteins with heat-induced misfolding (Fig. 5C).

**Figure 5.**
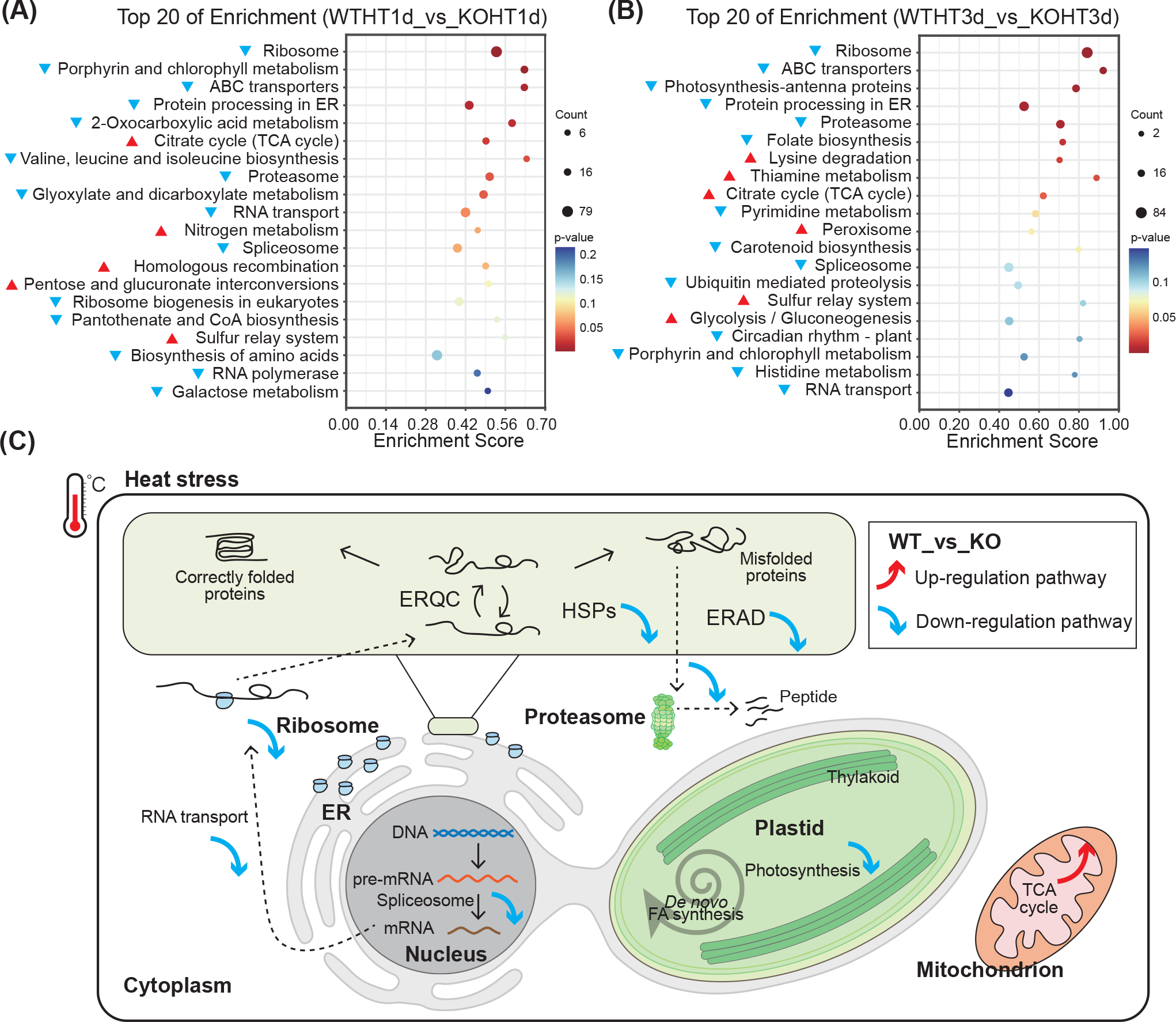
Transcriptome analysis of *ptELO5a* mutants and WT under heat stress (28°C). (A) (B) The enriched metabolic pathways of significantly differentially expressed genes between ptELO5a mutants and WT at 1 and 3 days of heat stress, and the top 20 metabolic pathways were selected based on significance. The red triangle indicates that the pathway was up-regulated in *ptELO5a* mutants compared to the WT, while blue triangle indicates down-regulation. (C) ER-related pathways have significant differences between *ptELO5a* mutants and WT in response to heat stress. ERQC, ER quality control; HSPs, heat shock proteins; ERAD, ER-associated degradation. Red arrow indicates that the pathway was up-regulated in the mutant strain, and blue arrows indicate that the pathway was down-regulated.

### The mutation in *ptELO5a* significantly altered the content and composition of fatty acids and glycerolipid in *P. tricornutum*

To reveal the direct effects of *ptELO5a* mutation on changes in lipid composition and related gene expression, fatty acids (FAs), lipid profiles and transcriptome of *ptELO5a* mutants and WT were compared at both exponential and stationary growth phase under standard culture conditions at 22℃. The knockout of *ptELO5a* evidently resulted in a significant reduction in 22:6, but had no effect on the accumulation of 20:5. Since most FAs were synthesized in chloroplasts, palmitic acid (16:0) and palmitoleic acid (16:1) accumulation in the *ptELO5a* mutants at the stationary phase were significantly lower than that in the WT (Fig. 6A). This also resulted in a significantly lower total FA content of 716 nmol mg^-1^ dry cell weight (dcw) in the *ptELO5a* mutants than 902 nmol mg^-1^ dcw in the WT during the stationary growth phase. Analysis of relative content profiles of FAs revealed significant decrease in the 16:0, 16:1 and 22:6 types of FAs in the *ptELO5a* mutants compared to WT at both growth phases. This decrease in FAs was balanced by the increase in myristic acid (14:0), 20:5 and lignoceric acid (24:0) (Fig. 6B).

**Figure 6.**
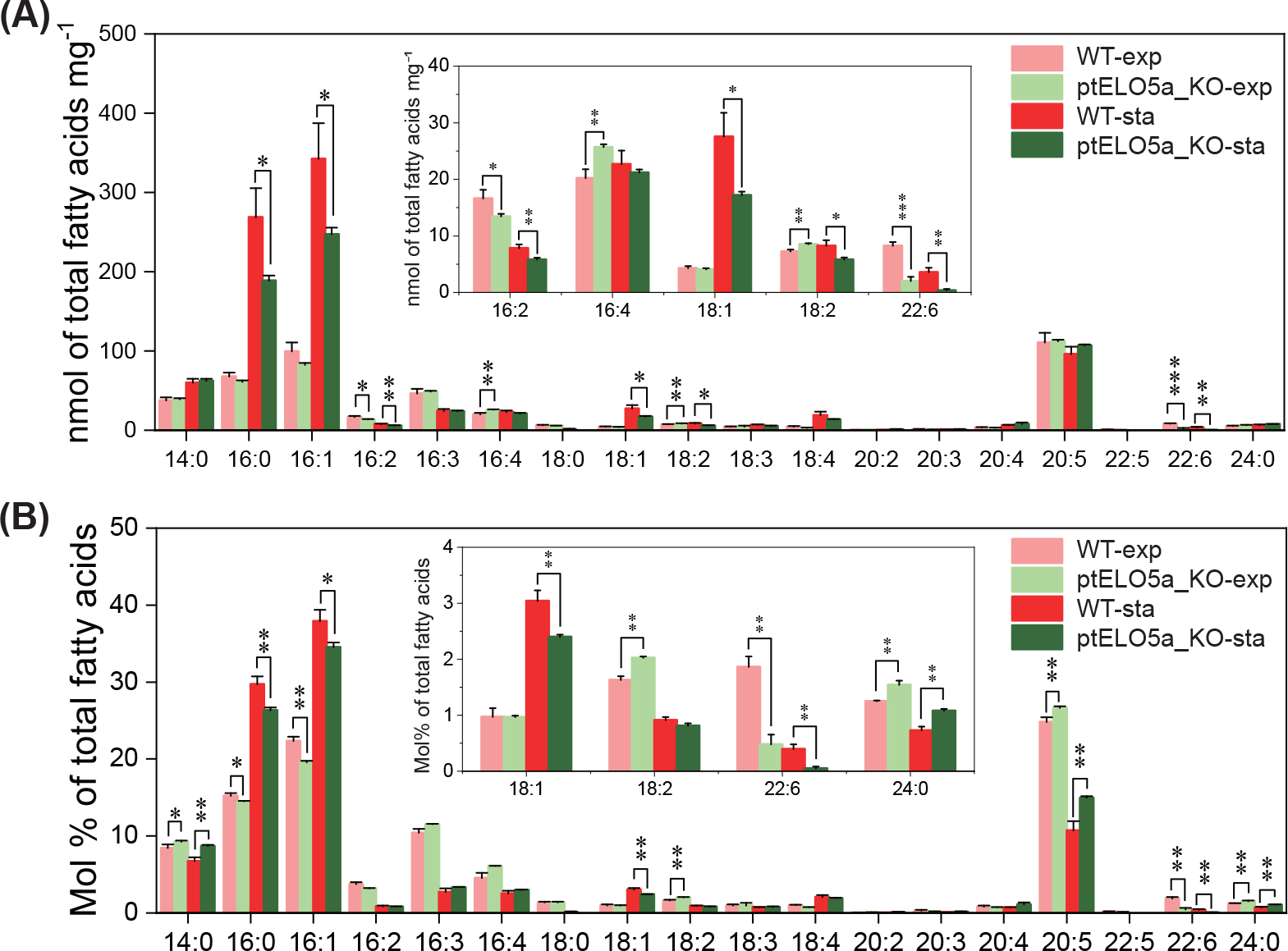
Total fatty acid content (A) and fatty acid composition (mol% of total fatty acids) (B) of WT and the *ptELO5a* mutants grown at exponential and stationary phases. Data and error bars are mean and standard deviation, respectively (n = 3). The asterisk indicates the significant difference (Student’s t-test, *, *P* < 0.05; **, *P* < 0.01; ***, *P* < 0.001) between the WT and *ptELO5a* mutants.

Glycerolipidomic analysis revealed that the inactivation of *ptELO5a* significantly affected the composition and distribution of different molecular species within each lipid class in the *ptELO5a* mutants (Fig. 7A). Compared to the WT, the disruption of *ptELO5a* gene in the mutants resulted in a significant increase in phospholipids in the exponential phase, including phosphatidylglycerol (PG, 52%, *P*<0.001), phosphatidylinositol (PI, 31%, *P*<0.01), phosphatidylethanolamine (PE, 47%, *P*<0.01) and phosphatidylcholine (PC, 21%, *P*<0.01). On the other hand, a significant reduction was observed in diacylglyceryl-hydroxymethyl-*N,N,N*-trimethyl-β-alanine (DGTA) as well as three abundant thylakoid glycolipids, including sulfoquinovosyldiacylglycerol (SQGD), monogalactosyldiacylglycerol (MGDG) and digalactosyldiacylglycerol (DGDG). In addition, the relative amount of TAG was also significantly lower in the *ptELO5a* mutants (*P*<0.05).

**Figure 7.**
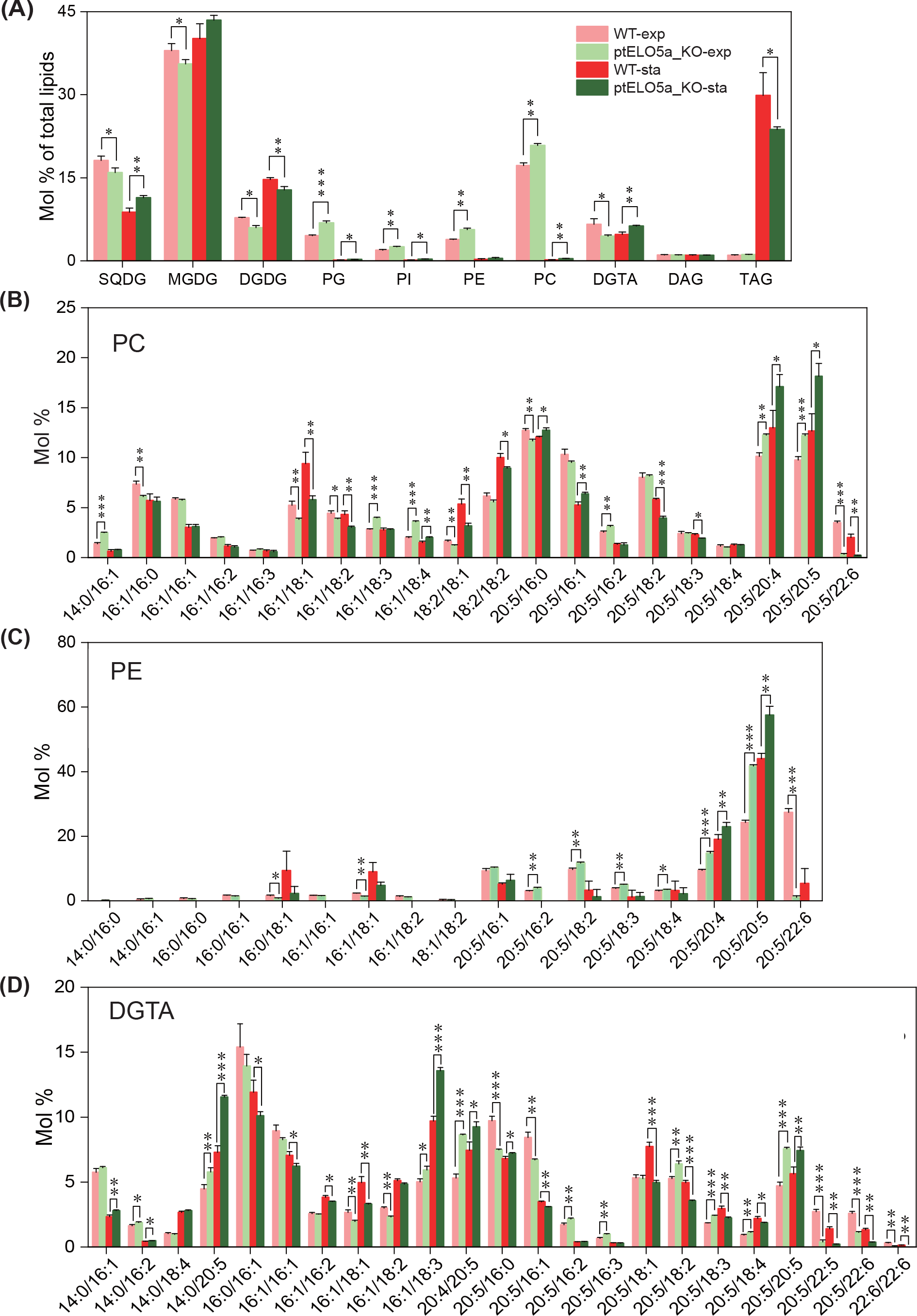
Comparative analysis of lipidome differences between WT and *ptELO5a* mutants grown at the exponential and stationary phases. (A) glycerolipids; (B) phosphatidylcholine (PC); (C) phosphatidylethanolamine (PE); (D) diacylglyceryl-hydroxymethyl-*N,N,N*-trimethyl-β-alanine (DGTA). Data and error bars are mean and standard deviation, respectively (n = 3). The asterisk indicates the significant difference (Student’s t-test, *, *P* < 0.05; **, *P* < 0.01; ***, *P* < 0.001) between the WT and *ptELO5a* mutants.

Analysis of the molecular composition of phospholipids showed that the knockout of *ptELO5a* directly affected the carbon chain elongation of 20:5 at the sn-2 position in phospholipid composition. This resulted in significantly higher 20:5 content in mutants, including 20:5/20:5 in PC and PE, as well as 14:0/20:5, 20:4/20:5 and 20:5/20:5 in DGTA. The knockout of *ptELO5a* also resulted in a significant reduction in the content of 22:6 and docosapentaenoic acid (22:5) in the three phospholipids, including 20:5/22:6 in PC and PE, and 20:5/22:5, 20:5/22:6 and 22:6/22:6 in DGTA (Fig. 7, B, C and D). The decrease in 22:6 content was mainly compensated by the increase in 20:5, while the total amount of 22:6 and 20:5 remained constant. Meanwhile, the excessive accumulation of 20:5 also caused the accumulation of the precursor fatty acids, including linolenic acid (18:3), parinaric acid (18:4) and arachidonic acid (20:4), for 20:5 synthesis in the *ptELO5a* mutants. Meanwhile, the relative contents of 16:0, 16:1 and oleic acid (18:1) significantly decreased in the mutants (Fig. S9).

The molecular composition of MGDG, DGDG and SQDG, which are main components of plastid membrane lipids in diatoms, were also affected in *ptELO5a* mutants (Fig. S10). The relative contents of 20:5/16:2 and 20:5/16:3 in MGDG accounted for more than 50%, and were significantly higher in the *ptELO5a* mutants. The DGDG composition of the mutants significantly reduced in the fraction containing 20:5 at the sn-1 position, whereas the proportions of all other fractions increased. In SQDG, the relative contents of 14:0/16:0 and 14:0/16:1 (the two most dominant fractions) were significantly higher in the mutants. Statistical analysis of the fatty acid composition of three glycolipids showed that the trend of the changes in fatty acids of DGDG was essentially opposite to that of MGDG, with a decrease in the content of PUFAs (Fig. S11, A and B). MGDG and SQDG had basically the same trend of the changes in the fatty acids, except for the opposite trend of changes in 20:5 (Fig. S11C). The relative composition of total fatty acids in the three glycolipids of mutant strains mainly showed a significantly higher percentage of 14:0 and 16:3, a significantly lower percentage of 16:0 and 16:1, and no significant difference in the percentage of 20:5 (Fig. S11D). A heat map was constructed to show the variations in the percentage of 20 major TAG species. The heat map showed no significant difference in the three major components with highest relative content, including 48:2 (16:0_16:1_16:1), 48:1 (16:0_16:0_16:1), and 48:3 (16:1_16:1_16:1). The percentage of TAG species containing 20:5 increased from 13.27% to 16.05% after mutation (Fig. S12).

### Suppression of central carbon metabolism in the *ptELO5a* mutants was the main cause of reduced TAG accumulation capacity

The production of lipids usually begins in the exponential phase and the accumulation of TAG occurs during the stationary phase in eukaryotic algae (Guschina and Harwood, 2006). However, neutral lipid content and triglyceride content revealed that the mutants were significantly weaker in accumulating TAG during stationary phase compared to WT (Fig. 8, A and B). In this study, mRNA-seq analysis was performed to understand how *ptELO5a* knockout affected the lipid accumulation in *P. tricornutum*. To eliminate the interference of other factors on gene expression, the transcriptome of the *ptELO5a* mutants and WT cells cultured under optimal growth conditions were compared at exponential and stationary phases. Enrichment analysis of KEGG pathways in significant DEGs showed that the differential pathways mainly included some pathways related to carbohydrate and lipid metabolism (Fig. 8C). Most DEGs of these pathways mostly related to carbohydrate metabolism (i.e., pyruvate metabolism, pentose phosphate pathway, and glycolysis/gluconeogenesis) are significantly suppressed in mutants in stationary phase (Fig. 8, D and E). In addition, the degradation of valine, leucine, and isoleucine (another important pathway that provides substantial substrates for lipid accumulation) was also inhibited in the mutants (Fig. S13).

**Figure 8.**
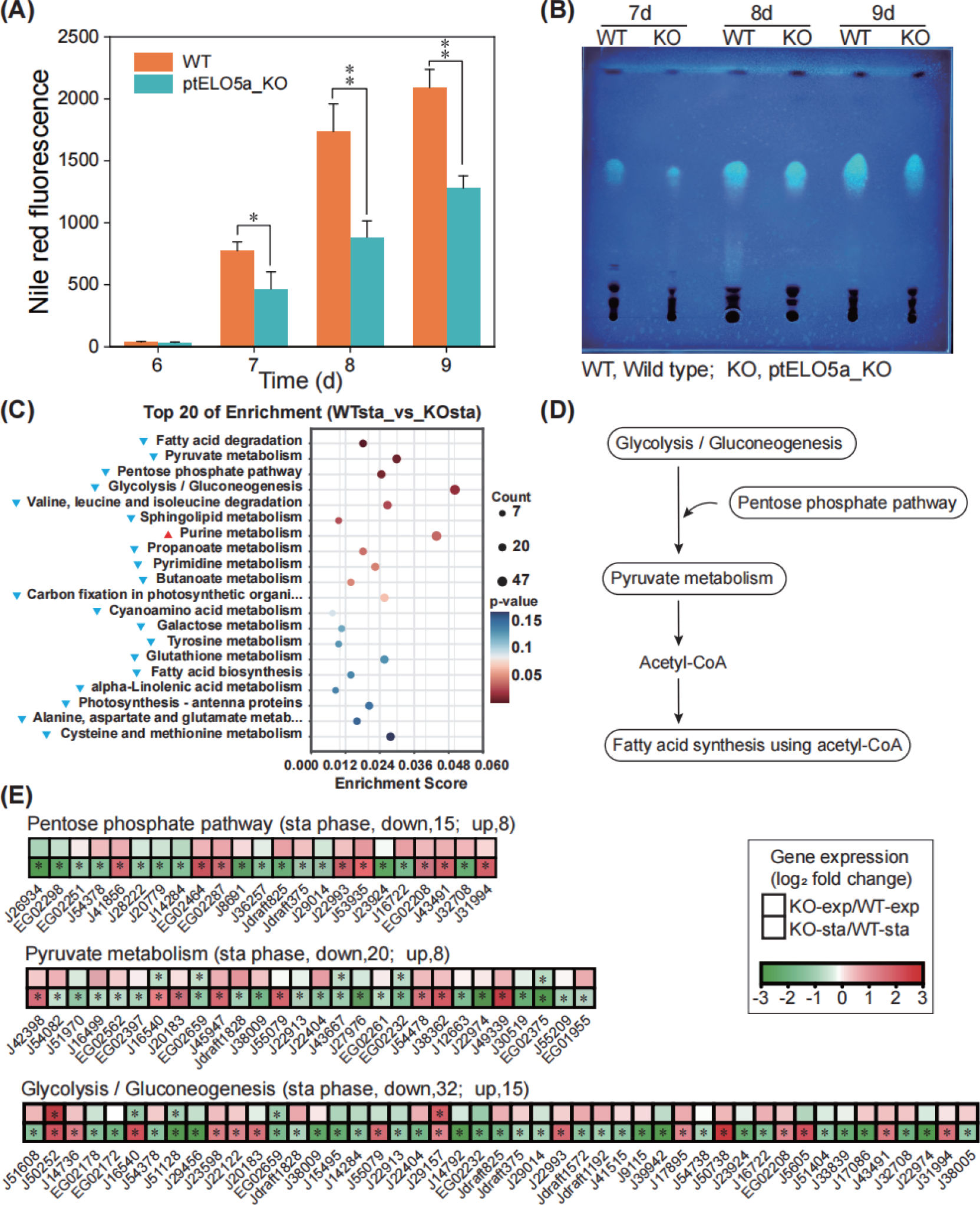
Impacts of *ptELO5a* knockout on lipid accumulation and transcriptome under standard culture conditions (22°C). (A) Nile red fluorescence intensity of *ptELO5a* mutant and WT cells grown for 6, 7, 8 and 9d, respectively. (B) A thin-layer chromatogram of total lipids from WT and *ptELO5a* mutants grown for 7, 8 and 9d respectively. Each lipid sample was extracted from ∼5×107 cells. TAGs were visualized with 0.01% (w/v) primuline reagent. (C) KEGG enrichment analysis of differentially expressed genes at stationary phase between the WT and *ptELO5a* mutants. Samples were collected on 8d. (D) A simplified model of the central carbon metabolic pathway responsible for lipid metabolism based on significant difference pathway analysis. (E) Major significant difference genes in central carbon metabolism (pentose phosphate pathway, pyruvate metabolism and glycolysis/ gluconeogenesis) between WT and *ptELO5a* mutants. Data are mean of log2(fold changes) (n=3) and are presented as heat maps with shades of red or green colors according to the scale bar. Fold changes were calculated as log2(FPKM (*ptELO5a* mutants) / FPKM (WT)). FPKM, absolute abundance of transcripts. *, |log2(fold change) | > 1.

The expression of some key lipid-related genes in the mutants were also down-regulated during stationary phase (Supplemental Dataset S4), especially the encoding genes of acetyl-CoA carboxylase (ACC1, Phatr3_J55209; ACC2, Phatr3_EG01955) and malonyl-CoA:ACP transacylase (MCAT, Phatr3_J37652), which are critical for the utilization of acetyl-coenzyme A for fatty acid synthesis and carbon chain elongation in plastids and cytoplasm. Acyl-CoA:diacylglycerol acyltransferase (DGAT2D, Phatr3_J43469) and phospholipid:diacylglycerol acyltransferase (PDAT, Phatr3_J8860), which are directly involved in TAG synthesis, were also significantly down-regulated in the mutants. In addition, the vast majority of genes in the TAG and fatty acid degradation pathways were also significantly down-regulated, with the exception of the encoding gene of acetoacetyl-CoA thiolase (Perox-AACT, Phatr3_J45947; Mito-AACT2, Phatr3_J28068), glyoxisomal malate dehydrogenase (Mito-PMDH, Phatr3_J42398) and citrate synthase (Mito-CSY, Phatr3_J30145).

### Complementation of *ptELO5a* mutants restored their heat tolerance and TAG accumulation capacity

To further verify that the changes in lipid content were caused by *ptELO5a* knockout rather than a second point mutation, complementary algal strains of *ptELO5a* (*ptELO5a-*Com) were constructed (Fig. 9A). To avoid knockout of backfill sequences, free knockout plasmid-removed knockout strains were obtained by successive zeocin-free passages (Fig. S14). The analysis of fatty acid composition revealed that the relative content of 22:6 in the *ptELO5a-*Com strains significantly elevated to the level similar to the wild type (Fig. 9B). Under heat stress, the growth of the *ptELO5a-*Com strains was basically the same as that of the wild type both grown in the cultures bubbled with filtered air and in the static culture (Fig. 9C; Fig. S15). In addition, the growth of the *ptELO5a* overexpression strains under heat stress was also consistent with that of the wild type (Fig. S16). The Nile Red-stained fluorescence and TLC separation assay further confirmed that the TAG accumulation pattern in the *ptELO5a-*Com strains and WT during stationary phase was also consistent (Fig. 9, D, E, F and G). These results confirmed that the phenotypes of the *ptELO5a* mutants were indeed caused by the inactivation of the *ptELO5a* gene. This finding excluded the possibility of second point mutation.

**Figure 9.**
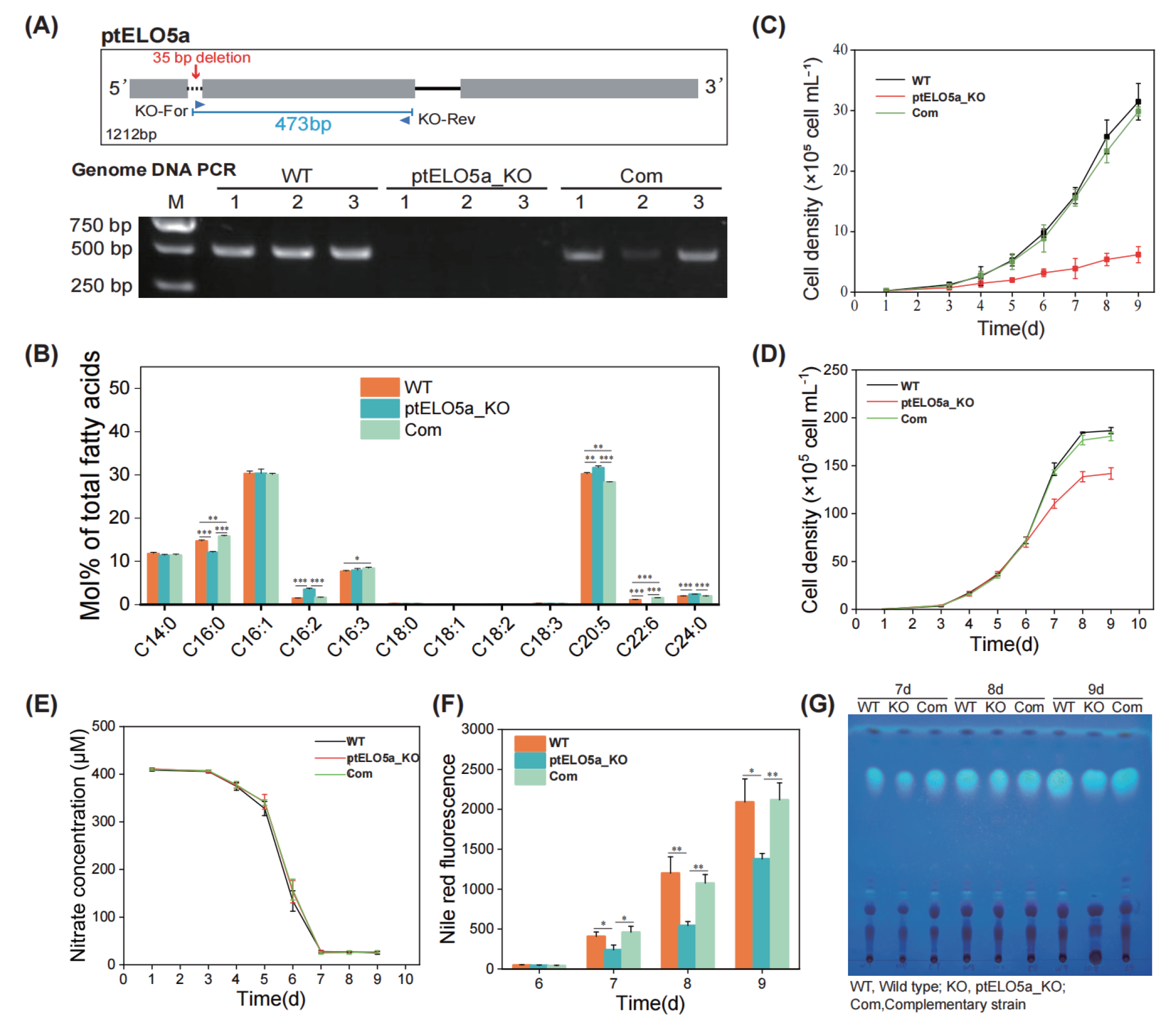
Phenotype analyses of *P. tricornutum* wild type (WT), *ptELO5a* mutants (ptELO5a_KO) and complementary strains (Com). (A) PCR of genomic DNA of the complementary strains using the ELO5aKO_colony-F/R primers. (B) Comparative analysis of fatty acid composition (mol% of total fatty acids) of wild type, *ptELO5a* mutants and complementary strains at exponential phase at 22℃. (C) Growth curves of wild type, *ptELO5a* mutants and complementary strains under air bubbling condition for 9 d at 28℃. (D) Growth curves of wild type, *ptELO5a* mutants and complementary strains under air bubbling condition for 9 d at 22℃. (E) Nitrate concentrations in the f/2 media for wild type, *ptELO5a* mutants and complementary strains for 9 d at 22℃. (F) Nile red fluorescence intensities of wild type, *ptELO5a* mutants and complementary strains grown for 9 d at 22℃. (G) A thin-layer chromatogram of total lipids from wild type, *ptELO5a* mutants and complementary strains grown for 7 d, 8 d and 9 d at 22℃. Each lipid sample was extracted from ∼5×107 cells. TAGs were visualized with 0.01% (w/v) primuline reagent. Data and error bars are mean and standard deviation, respectively (n = 3). The asterisk indicates the significant difference in two-by-two comparisons between the three algal strains (Student’s t-test, *, *P* < 0.05; **, *P* < 0.01; ***, *P* < 0.001).

## Discussion

### The role of long-carbon-chain elongase in lipid remodeling and environmental adaptation has been underestimated in marine phytoplankton

Both marine eukaryotic phytoplankton and terrestrial higher plants evolved from prokaryotic cyanobacteria due to the endosymbiosis during the long evolutionary process (Zimorski et al., 2014). Unlike the terrestrial higher plants that can only synthesize C16 or C18 PUFAs, many marine phytoplankton also have the ability to synthesize LC-PUFA (more than 18 carbon atoms) (Dolch and Maréchal, 2015). In general, the LC-PUFAs synthesis pathway requires multiple elongases and desaturases, which are mainly distributed in the cER or ER, and the synthesis of 20:5 from 18:1 can be categorized into the traditional Δ6 pathway or the alternative Δ8 pathway (Gong et al., 2014; Huang et al., 2023). And the synthesis from EPA to DHA requires two conserved steps: elongation of EPA to docosapentaenoic acid (DPA, 22:5 n-3) by ELO5, and then desaturation by Δ4 desaturase (DES4) (Ruiz-López et al., 2012).

In addition to its speculated role in regulating the phase transition temperature of membranes, EPA is also thought to be closely related to photosynthesis due to the high amount of EPA in glycolipids in marine phytoplankton (Abida et al., 2015). In comparison, DHA content is usually much lower than EPA and mainly found in phospholipids with unclear functions (Sayanova et al., 2017; Zulu et al., 2018). Recent extensive field investigations and lipid analysis in the ocean have revealed that the abundance of EPA in marine phytoplankton decreases with the increasing temperature, while DHA content first increases and then decreases with increasing temperature (Holm et al., 2022). This suggested that DHA is less sensitive to temperature than EPA. This discrepancy in temperature sensitivity indicates that DHA and EPA may have different physiological functions in phytoplankton. So far, the roles of DHA and carbon chain elongation, especially long-chain elongation enzymes, in the environmental adaptation of marine phytoplankton have been overlooked.

### The knockout of *ptELO5a* affects not only DHA synthesis but also lipid metabolism

Both overexpression and knockdown experiments indicate that PtELO5a is the dominant Δ5 elongases. Although the *ptELO5a* mutants did not have a reduced overall fatty acid synthesis capacity during the exponential phase, the percentage of glycolipids and phospholipids changed significantly (Fig. 7A). Phospholipids such as PG, PI, PE, and PC were significantly increased, whereas glycolipids such as MGDG, DGDG, and SQDG were significantly reduced. Transcript levels of the genes related to synthesis of phosphatidic acid (PA), diacylglycerol (DAG), and cytidine diphosphate diacylglycerol (CDP-DAG) in plastids and ER were upregulated in the mutants, including the genes encoding glycerol-3-phosphate acyltransferase, 1-acylglycerol-3-phosphate acyltransferase, and CDP-DAG synthase. The synthesis of polar head precursors in the cytoplasm as well as synthesis, desaturation, and polar head exchange of PE, PS and PC in the ER were up-regulated in the mutants. Moreover, the transcript levels of most of the genes involved in the synthesis of galactolipids and sulfolipids from DAG in plastids were significantly down-regulated in the mutants, which is likely the direct cause of the reduced levels of glycolipid synthesis (Supplemental Dataset S4).

Mutations in *ptELO5a* also resulted in a reduced ability to accumulate TAGs. This may affect changes in the ability of mutants to adapt to environmental conditions, as TAGs are the main storage lipids in photosynthetic organisms under stress (Yang et al., 2022). DHA is mainly distributed on phospholipids, which are low in content and not directly involved in TAG synthesis in wild-type *P. tricornutum* (Abida et al., 2015). Thus, there may be some indirect correlation between DHA content and the ability to accumulate TAGs. This was revealed by further analysis of the transcription of genes associated with lipid accumulation. Several important pathways involved in carbohydrate metabolism as well as the branched-chain amino acid degradation pathway are significantly repressed in the *ptELO5a* mutants, which greatly reduces the availability of carbon precursors (Fig. 8C; Fig. S13). The degradation products of plastid proteins and polar lipids have been shown to be the main carbon precursor resource for TAG (Ge et al., 2014; Levitan et al., 2015); The expressions of ACCs and MCAT, which are critical for the initiation of fatty acid *de novo* synthesis, were significantly reduced in the mutants, which might have been highly detrimental for the translation of acetyl-CoA in the overall lipid synthesis pathway (Zulu et al., 2018). Thus, insufficient supply of carbon precursors and down-regulation of key node genes may be the main reasons for the reduced accumulation capacity of TAGs in *ptELO5a* mutants.

### Disruption of *ptELO5* caused heat sensitivity in *P. tricornutum*

In *P. tricornutum*, DHA is mainly incorporated into the *sn*-2 position of glycerol backbone of PC, PE and DGTA. Esterification of PUFAs to membrane lipids is crucial to maintain the proper membrane fluidity, especially at low temperature (Valentine and Valentine, 2004). For instance, arachidonic acid (20:4)-containing MGDG might contribute to maintenance of chloroplast membrane fluidity at low temperatures in the green alga *Lobosphaera incisa* (Zorin et al., 2017).

In this study, *ptELO5a*-disrupted mutants were prepared to gain new insight into the physiological role of DHA in *P. tricornutum*. The growth of *ptELO5a* was normal at 22°C, whereas it was strongly impaired at 28°C. The limiting temperature for keeping membrane integrity in the wild type was 28.5°C (Cheong et al., 2021). Apparently, the altered membrane lipid composition in the *ptELO5a* mutants significantly reduced the tolerance temperature. Here, attempts were made to establish a link between the impaired growth and lipidomic changes. Lipidomic analysis showed that in the *ptELO5a* mutants, the most dramatic change in the proportion of DHA-containing glycerolipid molecular species was observed in PE 20:5/22:6, which decreased from 27.4% in WT to 1.11% in the mutants during the exponential phase, and from 5.35% in WT to undetectable level in the mutants in the stationary phase (Fig. 7). A significant reduction was also observed in other DHA-containing molecular species, including 20:5/22:6 of PC, and 20:5/22:6 and 22:6/22:6 of DGTA in the *ptELO5a* mutants. Furthermore, the impact of PE:PC ratio has been reported on the regulation of membrane fluidity in higher eukaryotic cells, such as insect and mammalian cells (Dawaliby et al., 2016). In this study, PE:PC ratio significantly increased from 0.22 in the WT to 0.27 in the *ptELO5a* mutants, indicating the change in the cellular membrane fluidity in *ptELO5a* mutants, especially at higher temperature. In marine cyanobacteria *Synechococcus*, fatty acid moieties of glycolipids were modified in response to temperature variation. When growth temperature decreased, the average acyl chain length of galactolipids decreased and the global proportion of unsaturated chains in the membranes strongly increased (Pittera et al., 2018). Compared to WT, the average length of the PC and PE *sn*-1 and *sn*-2 acyl chains significantly decreased in the *ptELO5a* mutants (Supplementary Table S4). This change may affect the functionality of extraplastidic membranes in the *ptELO5a* mutants, causing impaired cell growth at higher temperature of 28°C.

In addition, we also found that some metabolic pathways in the mutants were more affected under heat stress, mainly reflecting the close association with organelles composed of phospholipids (Fig. 5). The quality of the ER is considered critical to the stress response of algal cells, and it is also closely related to phospholipid composition (Yamaoka et al., 2019). Moreover, the rapid response of ER quality control (ERQC) and ER-associated degradation mechanism (ERAD) at high temperatures, as well as the rate of degradation of misfolded proteins in the ER system are critical for heat tolerance of algae (Chen et al., 2022). In addition, the gene expression

levels related to the majority of heat stress proteins (HSP) in the *ptELO5a* mutants were significantly lower than those in the WT, which may largely affect the thermotolerance of mutants (Ding et al., 2020). Maintenance of protein processing machinery and membrane structure is important for temperature acclimation and adaptation in marine diatoms (Liang et al., 2019). However, the expression of these genes was significantly suppressed in the mutants, which is clearly unfavorable for the mutants to cope with high temperature stress.

### The distinct physiological functions of EPA and DHA in marine phytoplankton

EPA and DHA may have distinct physiological functions in marine phytoplankton. Unlike EPA, which ends up localizing into the chloroplast membrane galactolipids (SQDG, MGDG and DGDG) after its synthesis in the ER, DHA is mainly distributed in PC, PE and DGTA, suggesting that DHA is important for the functionality of extraplastidic membranes. The changes observed in the ultrastructure of chloroplast in the *ptELO5a* cells might have been caused by the altered EPA composition in plastid membrane lipids. Alternatively, there may be a tight association between chloroplast membranes and extraplastidic membranes. Intercepting the synthesis of DHA, the original DHA-containing fraction was replaced by EPA, resulting in reduced adaptation to adversity in the mutants. The DHA synthesis of the constructed complementary strains was basically at the same level as that of the wild type in terms of temperature adaptability and neutral lipid synthesis. Thus, the role of small amounts of DHA in regulating the physiology and cellular structure of phytoplankton cannot be ignored (Fig. 10). DHA synthesis is at the end of the entire LC-PUFAs synthesis pathway and is often overlooked in studies, but it is important in the regulation of overall lipid composition and stress adaptation in diatoms.

**Figure 10.**
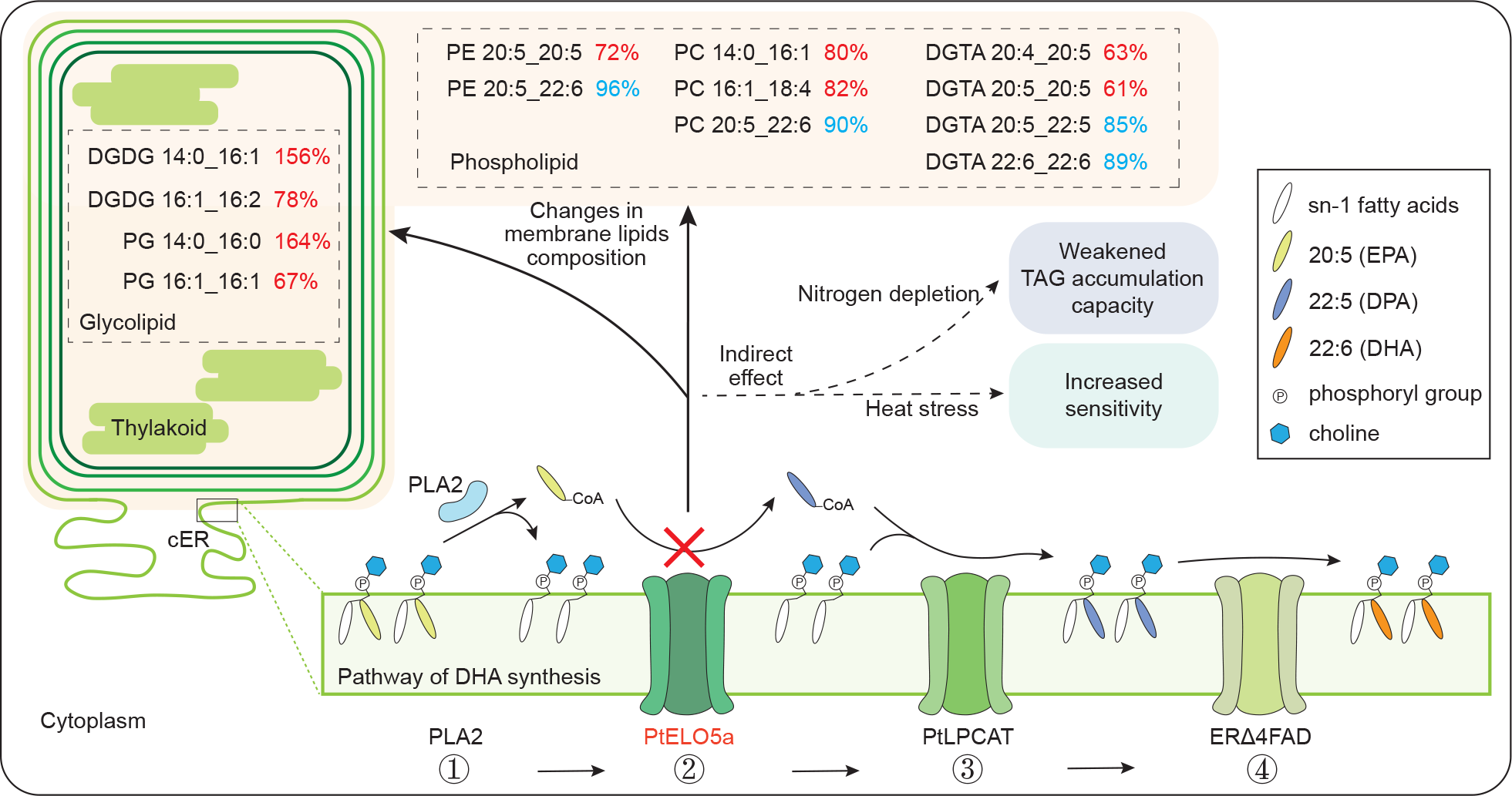
Schematic representation of the PtELO5a-mediated lipid metabolism pathway in *P. tricornutum*. The continued catalytic synthesis of DHA from EPA, which is abundant on phospholipids, occurs in four main steps. Step 1, Phospholipase A2 (PLA2) catalyzes sn-2 of phosphatidylcholine to form lysophosphatidyl- choline (LPC) and 20:5-CoA. Step 2, PtELO5a-catalyzed carbon chain elongation of 20:5-CoA to 22:5-CoA. Step 3, Acyl-CoA:lysophosphatidylcholine (PtLPCAT) recombines free 22:5-CoA with LPC in cER (You et al., 2023). Step 4, ER fatty acid desaturases 4 (ERΔ4FAD) catalyzes 22:5 to 22:6 in PC sn-2 in cER (Huang et al., 2023). By interrupting *ptELO5a* in this synthesis pathway, DHA synthesis is blocked, leading to alterations in the composition of both phospholipids and glycolipids, which in turn affects the response of the mutants to stress. Demonstrates glyceride fractions with a relative change greater than 60%. The relative content of the fractions is shown in red for elevated values in the mutants compared to the wild type and in blue for decreased values.

Under the scenario of future global warming, understanding the physiological functions and biosynthetic pathways of LC-PUFAs in marine phytoplankton not only helps to understand the changes and distribution of phytoplankton communities, but also has important significance for predicting the resource distribution of natural LC-PUFAs. High contents of EPA and DHA have been reported in many phytoplankton, not only in the polar marine environments, but also in subtropical and tropical marine environments with higher temperatures (Hixson and Arts, 2016; Zulu et al., 2018). Although phytoplankton can actively adjust their adaptive strategies under thermal variations, ocean warming will affect their physiological state and cellular composition, leading to changes in their eco-regional distribution (Liang et al., 2019; Li et al., 2023). The availability of LC-PUFAs and the nutrient quality of planktonic food web components will reduce under ocean warming, which will also significantly reduce the efficiency of human access to LC-PUFAs (Lau et al., 2021). DHA, like other LC-PUFAs, is also susceptible to oxidation under high-temperature stress, so algae tend to maintain higher levels of LC-PUFAs at lower temperatures (Jiang and Gao, 2004). However, the differences between DHA and EPA in the physiological regulation of algal cells should not be neglected. Our study suggests that those diatoms have maintained a strategy of elongating fatty acid from 20 to 22 carbon atoms over long periods of evolution, which to a certain extent enhances their ability to cope with high temperature. The synthesis of DHA and the elongation process of PUFA carbon atoms from 20 to 22 may be more important for them in the context of future climate change. Those diatoms with this elongation enzyme or DHA biosynthesis capability may have a competitive advantage in adapting to global ocean warming, which needs to be further revealed in the future.

## Funding

This work was supported by the grant from the National Natural Science Foundation of China (No. 31961133008 to YG; No. 32170108 to HJ) and the Science and Technology Innovation 2025 Major Project of Ningbo City (Grant No. 2022Z189) and Ningbo Public Welfare Research Program Project (No. 2023S040) to HJ. A. Amato, JJ and EM were also supported by Agence Nationale de la Recherche (ANR-10-LABEX-04 GRAL Labex, Grenoble Alliance for Integrated Structural Cell Biology; ANR-11-BTBR-0008 Océanomics; IDEX UGA CDP Glyco@Alps; Institut Carnot 3BCAR).

## Author contributions

J.Z., S.L. and W.C. performed experiments; X.X. and XP.W. provided technical assistance; XW.W., J.H. and H.H. analyzed the data. J.J., A. Amato and E.M. provided specific expertise in glycerolipid analyses; A. Allen contributed to the multiplexed CRISPR/Cas9 vectors and methods; Y.G. and H.J. planed and designed the research. J.Z. wrote the manuscript and Y.G. and H.J. revised the manuscript.

## Supporting information

Additional Supporting Information may be found online in the Supporting Information section at the end of the article.

## Competing interests

The authors declare that they have no conflict of interest.

## Accession numbers

Sequence from *Phaeodactylum tricornutum* genome and data used for phylogenetic tree reconstructions can be found in the *P. tricornutum* and the GenBank data library under the following accession numbers: Phtra3_J9255; Phatr3_J34485; KOO22560; EOD07354; CEF39061; AAL37626; ADD51571; BAI40363; ADN94475; ADN94476; GFH52305; AEA07666; BAO27787; OEU22482; ACK99719; AAT85662; ADE06662; AQX92136; AAV67797; AAW70157; AFU3574; ACR53359; AHG94993; GAY03028; AAV67799; GFH61049; KAG8470089; AFF27584; GAX16207; OEU09261; KAG7344361; AAV67798; AAV33630; VEU39922; ACR53360; AAY15135; AAV67800; KAI8329854; KAF8984070; GJJ77693; GAX16936; KAI2494228; OEU22771; KAI7818337; KAF9207334; KAF9933269; GKZ01156; KAG7344733; CAH0379219; KAI8371721; CAB9508581; CAB9525579; KAI9274193; CAE7610831; GMI28703; GMI31968; GMI26318; QDZ23655; CAJ1934847; KAI2494718; GKY98428; KAG7370759; GMI60985; KAJ1623087; CAB9512426; KAK1738254; GMI30663; GMH53578; GMH56579; GMH70427. Transcriptome sequence data are available at NCBI under BioProject accession PRJNA1055175 (https://www.ncbi.nlm.nih.gov/bioproject/?term=PRJNA1055175).

## Supplemental data

The following materials are available in the online version of this article.

**Supplemental Figure S1.** Analysis of the PtELO5a sequence.

**Supplemental Figure S2.** Analysis of the PtELO5b sequence.

**Supplemental Figure S3.** Subcellular localization of PtELO5a

**Supplemental Figure S4.** Construction of *ptELO5a/b*-OE strains.

**Supplemental Figure S5.** Fatty acid composition of wild type and *ptELO5a/b*-OE strains.

**Supplemental Figure S6.** Proportion of cells with abnormal morphology of *ptELO5a* mutants under heat stress.

**Supplemental Figure S7.** Enrichment analysis of differential metabolic pathways in heat stress experiments.

**Supplemental Figure S8.** Analysis of the differential expression of 9 heat shock protein (HSP) genes in the wild type and ptELO5a mutants in heat stress experiments.

**Supplemental Figure S9.** Comparative analysis of fatty acid composition of phospholipid between wild type and the *ptELO5a* mutants.

**Supplemental Figure S10.** Comparative analysis of glycolipid differences between wild type and *ptELO5a* mutants.

**Supplemental Figure S11.** Comparative analysis of fatty acid composition of glycolipid between wild type and the *ptELO5a* mutants.

**Supplemental Figure S12.** The disruption of *ptELO5a* affects molecular composition of triacylglycerol.

**Supplemental Figure S13.** Analysis of transcriptional differences in the branched-chain amino acid degradation pathway in the *ptELO5a* mutants.

**Supplemental Figure S14.** Selection of non-transgenic *ptELO5a* mutants.

**Supplemental Figure S15.** Growth of *P. tricornutum* wild type, the *ptELO5a* mutants and complementary strains in the static culture for 10 d under heat stress.

**Supplemental Figure S16.** Growth of *P. tricornutum* wild type and *ptELO5a* overexpression strains in the static culture for 10 d under heat stress.

**Supplemental Table S1.** List of plasmids used in this study.

**Supplemental Table S2.** List of oligonucleotides/primers used in this study.

**Supplemental Table S3.** Distribution of differential genes in each comparison combination.

**Supplemental Table S4.** Relative changes in fatty acids esterified at two stereospecific (sn) positions of glyceride in ptELO5a mutants and wild type.

**Supplemental Dataset S1.** Elongase sequence.

**Supplemental Dataset S2.** RNA-seq data.

**Supplemental Dataset S3.** Lipidomic datasets.

**Supplemental Dataset S4.** Lipid-associated RNA-seq analysis.

**Supplemental Dataset S5.** Summary of statistical analyses.

